# AI-driven high-throughput automation of behavioral analysis and dual-channel wireless optogenetics for multiplexing brain dynamics

**DOI:** 10.1101/2021.09.23.461279

**Authors:** Woo Seok Kim, Jianfeng Liu, Qinbo Li, Sungcheol Hong, Kezhuo Qi, Rahul Cherukuri, Byung-Jun Yoon, Justin Moscarello, Yoonsuck Choe, Stephen Maren, Sung Il Park

## Abstract

Advances in behavioral optogenetics are limited by the absence of high-throughput pipelines for the automated analysis of behavior in freely behaving animals. Although a variety of AI algorithms has been proposed that enable automation of behavioral analysis, existing methods are generally low-throughput. In addition, optogenetic manipulation of neural circuits typically requires physical tethers to light sources, which limits the number of brain areas that can be targeted and thus constrains behavioral testing. Here, we develop a new wireless platform that combines a battery-free dual-channel optogenetic implant with an AI algorithm for high-throughput behavioral analysis. In our platform, a customized AI algorithm detected and quantified freezing behavior of rats that had undergone fear conditioning. Notably, our platform allows up to four enclosures to be monitored simultaneously. Wireless dual-channel optogenetic devices were implanted in the basolateral amygdala (BLA) to permit independent modulation of BLA principal neurons (red light, AAV-CaMKII-JAWS) or BLA interneurons (blue light, AAV-mDlx-ChR2) in freely behaving animals. *In vivo* validation with behaving rats demonstrates the utility of the telemetry system for large-scale optogenetic studies.

**Significance:** AI algorithms can enable automation of behavioral analysis and thereby facilitate the progress on behavioral optogenetics. Successful integration of advanced wireless dual-channel optoelectronic devices with biological systems can also yield new tools and techniques for neuroscience research, particularly in the context of techniques for optogenetics. Here, we propose a new approach that combines an advanced AI algorithm with a low power wireless telemetry system, yielding powerful capabilities in the understanding of brain functions and the evaluation of the behavioral consequences of neural circuit manipulations. *In vivo* studies using optimized systems demonstrate high-throughput automation of behavioral manipulation and analysis via AI-powered wireless telemetry.

## Introduction

Optogenetics has transformed our understanding of brain dynamics, in particular how neural circuits control sensory perception, cognitive processes, and behavior(1, 2). However, utilizing this powerful genetic tool relies on the ability to quantitatively evaluate the behavioral consequences of neural circuit manipulations(3, 4). Unfortunately, the absence of high-throughput pipelines for the analysis of behavior in untethered, freely moving animals has limited progress. Recently, deep convolutional neural networks (DCNNs) have been developed that enable computer vision-based quantitative analysis of behavior(5, 6). These methods involve Mask R-CNN, DeepLab, and DeepLabCut(7–9).

For example, Mask R-CNN allows for the detection of multiple objects in real-time and accurate segmentation results for each instance(7, 10). DeepLab has been shown to excel in various challenging segmentation tasks by employing DCNNs and fully connected conditional random fields (CRFs) for semantic image segmentation(11). Another example is DeepLabCut, which detects user-defined body parts of humans or animals in video recordings and accurately estimates the pose of those parts without relying on explicit markers. These technologies enable real-time quantitative behavioral analysis in behavioral neuroscience studies. However, these methods still suffer from a key limitation in their ability to be multiplexed so that multiple animals in multiple cages can be studied in parallel.

Another complication of optogenetics is a lack of switching mechanism that enables the modulation of neuronal activity at multiple spatial scales in real-time. To systematically study brain mechanisms underlying a particular behavior or cognitive process, it is important to manipulate neuronal activity broadly across brain structures(2, 12, 13). This can be accomplished by dual configurations that use two fiber optics, each of which interfaces with a target brain region, to modulate neural activity in targeted brain regions(14–16). Although this approach has some utility, physical tethers to light sources limit the range of manipulations and environments that can be used in behavioral studies(17).

We have recently demonstrated the utility of wireless technologies to overcome these limitations(18, 19). Importantly, our implementation of this technology uses low-power devices that do not require batteries, further reducing the weight and profile of the implants. Here, we further develop this technology and introduce a dual-channel optogenetic device that can be paired with a high-throughput, AI-driven behavioral analysis platform for optogenetic studies in multiple, freely behaving animals(20). The wireless telemetry system consists of 1) an AI algorithm for postural detection and automated behavioral analysis in multiple animals, 2) a miniaturized implantable dual-channel optogenetic brain implant, and 3) a low-power wireless transmission (TX) system for simultaneous control of multiple animal enclosures.

A customized Al algorithm (DeepLabCut 2.1.10.4 and detailed package information in ***SI Appendix*, Table S1**) detects freezing behavior, and miniaturized battery-free wireless dual-channel optogenetic brain implants permit independent/simultaneous control of channels or brain regions in a freely behaving animal. The low-power wireless power TX system paired with advanced radio frequency power control strategies allows for simultaneous control of multiple animal enclosures (up to 4). Together, the AI-assisted implantable wireless telemetry system enables high-throughput analysis of animal behaviors relevant to post-traumatic stress disorders (PTSD) and anxiety. *In vivo* validation of the system was conducted in experiments in which freezing behavior was disrupted by optical inhibition or excitation of basolateral amygdala (BLA) principal neurons or interneurons, respectively.

## Results

An overview of the proposed wireless telemetry system is depicted in **Fig. 1**. This system integrates an advanced AI algorithm with a low-power wireless TX system to enable high-throughput automation of behavioral control and analysis (**Fig. 1*A***). In this system, a webcam monitors animals in 4 enclosures and acquires and transmits video (25 fps) of animals to a PC in real-time. Next, the AI algorithm analyzes animal behavior and assesses freezing behavior after the TX system initiates optogenetic stimulation (such as optogenetic stimulation of BLA neurons that regulates conditioned freezing) (**Fig. 1*B***). A dual-coil antenna is installed in each cage and the wireless power TX system manages power delivery to the enclosures. **Fig. 1*C*** shows the simultaneous operation of wireless subdermal implants in four different animals in independent enclosures. A miniaturized dual-channel wireless brain implant consists of an energy harvester and a dual-probe. It harvests radio frequency (RF) energy from the wireless power TX system and converts RF energy into optical energy to illuminate targeted brain regions (**Fig. 1*D***). When implanted, it allows for simultaneous and independent control of two brain regions or cellular targets within a brain region in a freely behaving animal. Each of these processes has been implemented in the experiments described below.

**Fig. 1.**
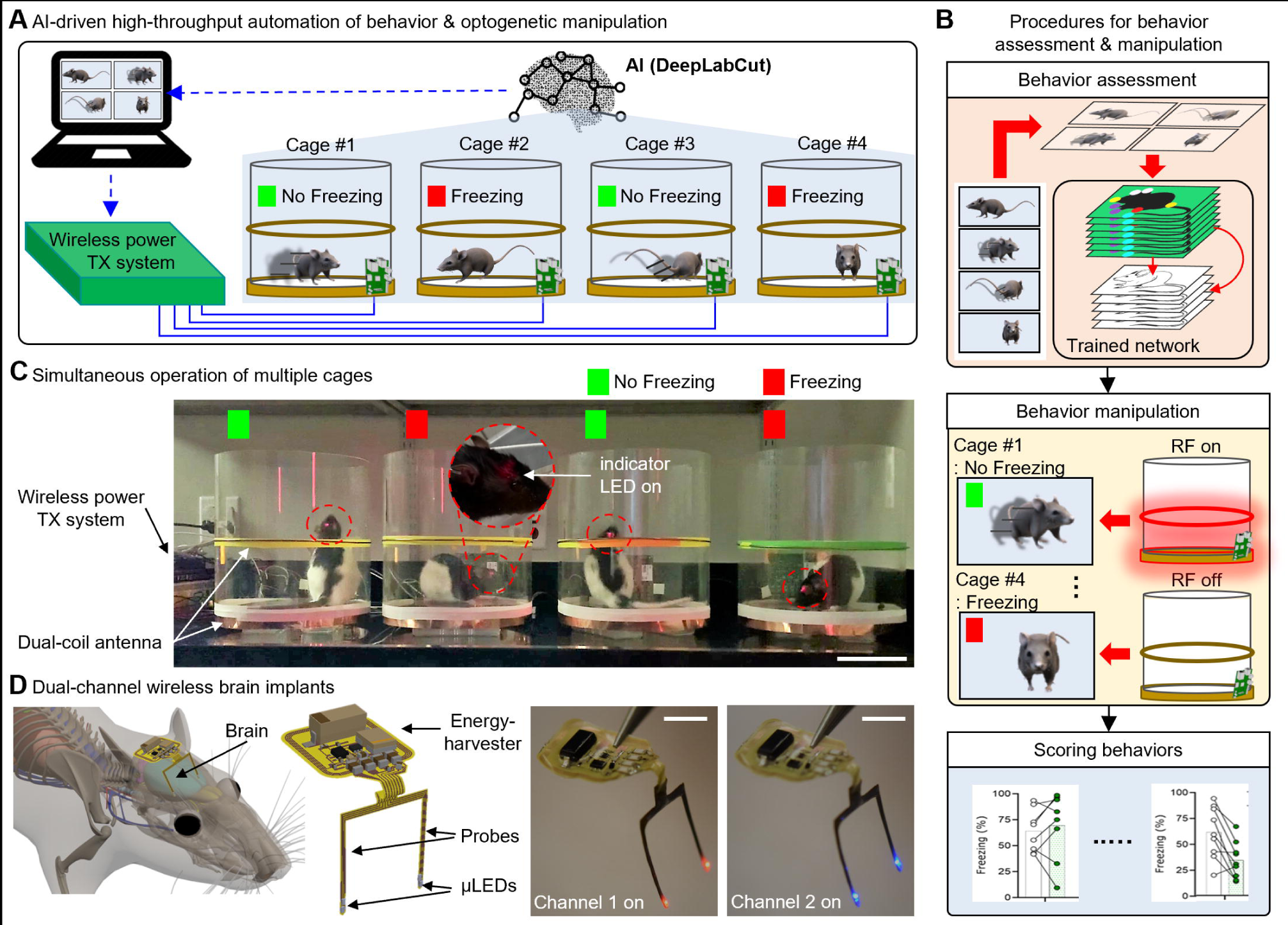
Overview of an AI-assisted implantable wireless telemetry platform for the study of brain dynamics and behavior. (*A*) Schematic illustration of the proposed AI wireless system. Here, the AI platform monitors animal behaviors, performs behavioral manipulation and assessment, and manages radio frequency power delivery to cages. (*B*) Procedures for behavior assessment and optogenetic manipulation. The AI algorithm analyzes framed images from the camera in realtime, assesses behavioral manipulation initiated by the TX system, and yields scoring results for observed freezing behaviors in the 4 cages. (*C*) Image of a high-throughput wireless power TX system and animals with the brain implants. Red dotted circles highlight wireless operation in freely behaving animals; scale bar 10 cm. (*D*) Schematic illustration of a dual-channel brain implant and images of wireless operation; scale bar 500 μm.

### AI-driven high-throughput automation of behavioral analysis

We have developed an AI-driven approach to analyze freezing behavior based on the position of defined regions of the rat’s body position. Our approach eliminates the constraint that the scene needs to be static, and thus generalizes well in real-world environments. Here, we trained the body part detection network to detect nine body parts in total: nose (N), two ears (LE and RE), two forelimbs (LF and RF), two hindlimbs (LH and RH), tail root (TR), and tail end (TE) because not all body parts of the rodent are always visible to the camera. **Fig. 2*A*** describes the overall workflow of our approach. Here, we used DeepLabCut proposed by Mathis et al. to detect pre-defined body parts(9). For behavior analysis/assessment of multiple animals, we first split the video and analyzed the freezing behavior independently, and then combine the results and videos at the end.

**Fig. 2.**
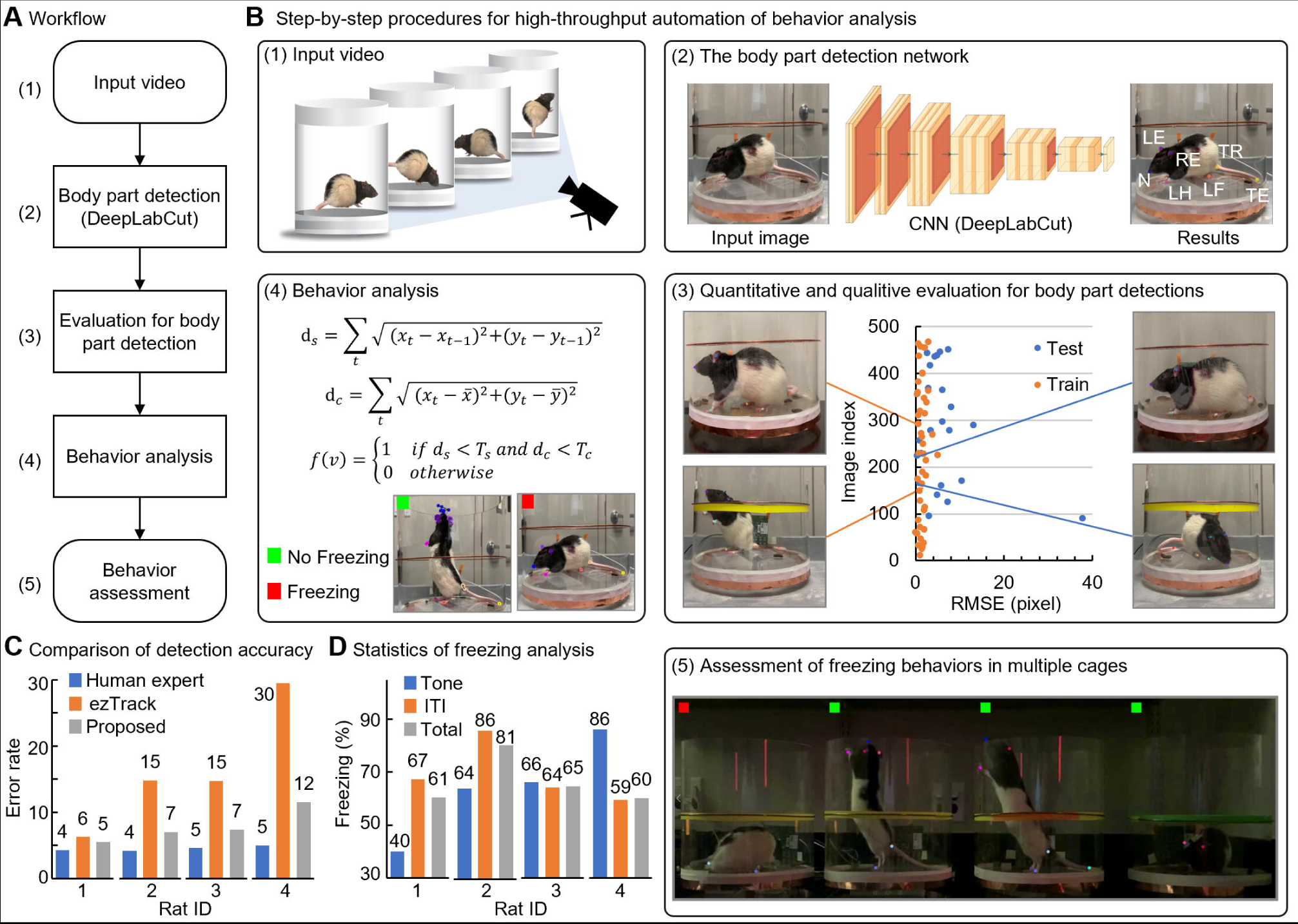
AI-enhanced high-throughput automation of behavior analysis. (*A*) A block diagram of workflows (*B*) Illustration of step-by-step procedures of proposed AI platform. The RMSE (Root Mean Square Error) for the training set and test set are 1.79 pixels and 8.32 pixels respectively; N, nose; LE & RE, two ears; LF & RF, two forelimbs; LH & RH, two hindlimbs; TR, tail root; TE, tail end;, the accumulated distance moved over;, the distance from the center point over;, the freeze detection results for video;, cutoff thresholds, 0.5 and 0.7, respectively. (*C*) Comparison of error rates with other methods (ezTrack and human observer). Here, we calculate the errors by human experts (n=3). (*D*) Statistics for freezing analysis in three different conditions: Tone (10 s 5 times), ITI (30 s 4 times), and Total (Tone ITI one 30 s post).

Step-by-step procedures for automation of behavior analysis appear in **Fig. 2*B***. First, we manually label 475 frames out of 5 videos captured in the environment described in **Fig. 2*B*(1)**. Next, we train our network for 300k iterations using the ResNet-50(21) (**Fig. 2*B*(2)**). To evaluate our detection network, we randomly split the labeled dataset into a training set with 95% of the frames and a test set with 5% of the frames. Here, we estimate the RMSE (Root Mean Square Error) between our results and human labels for quantitative evaluation. The RMSEs for the training set and the test set are 1.79 pixels and 8.32 pixels respectively. It is followed by the quantitative and qualitative evaluation of our body part detection network (**Fig. 2*C*(3)**). The detection network outputs three values for each body part: 1) x coordinate, 2) y coordinate, and 3) the likelihood of the detection. According to the defined thresholds *T_s_* and *T_c_*, the behavior analysis module can determine whether the subject’s status in the selected frame is freezing or not. Specifically, we set the time period *T* as 1 second in this setup: cutoff threshold *T_s_* as 0.5, and finally *T_c_* as 0.7. Finally, these processes yield the statistics of behavioral analysis in multiple cages where animals freely behave (**Fig. 2*B*(4) and (5)**). ***SI Appendix*, Movie S1.** provides visual evidence of the high-throughput setting.

To demonstrate the efficacy of the proposed behavior analysis method, we compared our method with ezTrack(22). Here, we capture four videos that do not overlap with the five videos used to train the body part detection network. As shown in **Fig. 2*C***, our approach outperformed ezTrack. The statistics of the analysis results of the developed software under three conditions revealed that the subjects are freezing for 64 % of the time with tone, and 69.2 % for ITI (**Fig. 2*D***). The result of unconditioned animals is shown in ***SI Appendix*, Fig. S1**. Empirically, we found that more than five body parts are usually visible to the camera, and at least two body parts are always visible. Compared with the conventional method(9) that uses four body parts in an overhead view, our approach (screen division in the side view) yielded an improved error rate.

### Dual-channel wireless optogenetic device

The dual-channel wireless brain implant employs a switching mechanism enabled by a Reed switch and a parallel capacitor(23, 24). A pulse wave transmitted from the TX system can discharge a capacitor, C1, connected in parallel with a reed switch, through a resistor, R1 (**Fig. 3*A***), and thereby turns off NMOS1 and activates PMOS1 & NMOS2. This allows for independent control of each channel. Here, we define the threshold time for switching and it is determined by resistance (R1) and capacitance (C1). A state diagram illustrates the switching mechanism in detail (**Fig. 3*A***). A pulse width shorter than the threshold does not provide enough time for C1 to discharge through R1, and NMOS1 remains activated (Ch1; red). When a longer pulse than the threshold is transmitted, C1 can fully discharge through R1. This would bring the voltage level to zero (ground) and consequently turns on PMOS1 and NMOS2. This results in the activation of Ch2 (blue).

**Fig. 3.**
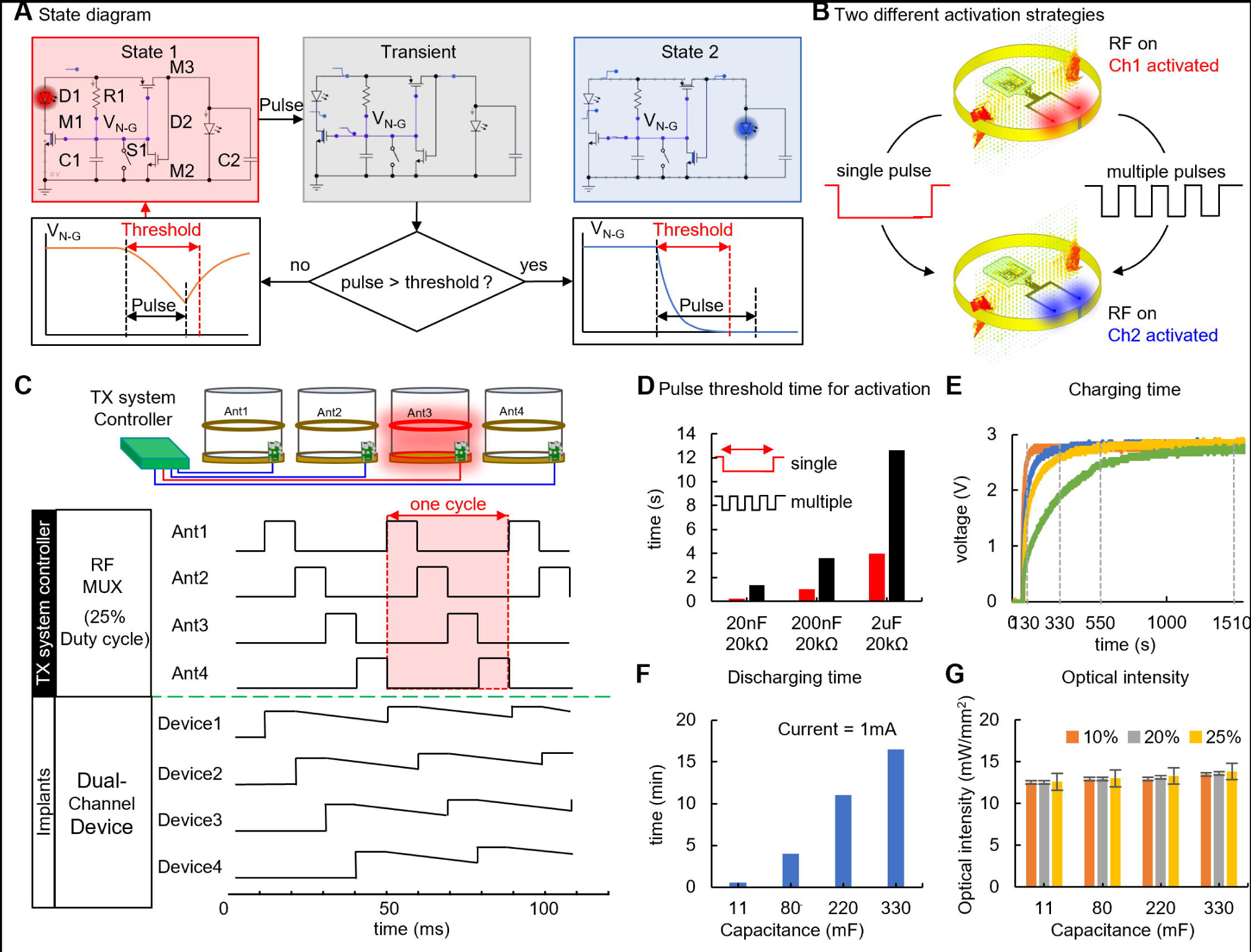
Characteristics of a high-throughput wireless power TX system. (*A*) State diagram of channel switching operation. Here, we capture transient responses at each node when a pulse signal shorter or longer than the threshold for switching, respectively, is applied to the device. M1, p-mos; M2-3, n-mos; R1, resistor; C1-2, capacitors; D1, red μLED; D2, blue μLED; S1, reed switch. (*B*) Illustration of two different strategies for channel switching. (*C*) Illustration of supercapacitor assisted time division multiplexing scheme and representative waveforms at each cage and implant. (*D*) Plot of threshold for switching as a function of time constant. (*E*) Plot of charging time as a function of capacitance. (*F*) Plot of discharging time as a function of capacitance. (*G*) Optical intensity vs. capacitance as a function of duty cycle.

The actuation mechanism offers two different strategies for switching: a single pulse and multiple pulses (**Fig. 3*B*** and ***SI Appendix*, Movie S2**). For a combination of 10 μF and 20 kΩ, a pulse width needs to be longer than 1 second for the activation of Ch2. This will ensure that charges accumulated in C1 can fully discharge through R1. This suggests that channel switching can be made if a voltage at V_N-G_ is pulled down to zero-level for a few seconds within the intervals. For example, multiple (e.g., 5) pulses (on; 500 ms & off; 500 ms) can activate Ch2 and it takes 5 seconds for switching. This indicates that the time constant (threshold) for charging is greater than that for discharging. That is, C1 discharges faster than it charges and thereby net charge would be zero after a few cycles. The threshold for activation or switching is tunable and one can use a different combination of resistors and capacitors to increase or decrease the threshold time (**Fig. 3*C*** and ***SI Appendix*, Fig. S2-4**). This would avoid unwanted channel activation or switching associated with environmental noise by a combination of large RC values (e.g., it would require a long pulse to switch a channel and thereby make it less sensitive to noise).

The wireless telemetry system enables simultaneous activation of implants in multiple cages (up to 4). Here, we use a time-division multiplexing strategy with the TX system to deliver RF power to each cage in a programmed manner. Although RF power is not available to all cages at all times, a wireless brain implant can store RF power for use later in a supercapacitor while supplying power to the circuits. This ensures continuous operation of brain implants regardless of duty cycle. The characteristics of this supercapacitor, including charging/discharging time and optical intensity during operation, are summarized in **Fig. 3*D-F*** and ***SI Appendix*, Fig. S5**. A supercapacitor with the capacitance of 80 mF (4.8 mm in diameter) requires 130 seconds for operation while 330 mF (6.8 mm in diameter) requires 1510 seconds (**Fig. 3*D***). Discharging time depends on load: here, LEDs or their power consumption. Measurement results revealed that a supercapacitor with capacitance ranging from 11 to 330 mF can supply a constant current of 1 mA for 1, 4, 10, and 16 min, respectively (**Fig. 3*E* and *F***). This will ensure an optical intensity over 10 mW/mm^2^, enough for activation of light-sensitive opsins when a brain implant does not receive RF power from the TX system during an off-cycle of the TX system. Measurements of optical intensity showed that the wireless TX system allows for simultaneous activation of multiple cages (**Fig. 3*G*** and ***SI Appendix*, Fig. S6**). Also, thermal assessment of the dual-channel implants demonstrated maximal temperature increases (~0.6 °C; ***SI Appendix*, Fig. S7**).

### Wireless optogenetic control of fear retrieval by the TX system

We then examined whether the TX system could function effectively to deliver wireless optostimulation in behaving animals. To explore this, we employed a widely used behavioral task; auditory fear conditioning(25). Freezing behavior was used as an index of fear in this behavioral task. We targeted the BLA, a brain region that is critical for the expression of conditioned freezing behavior in this task, to validate the TX system. We injected AAV8-CaMKII-Jaws-GFP or its control AAV8-CaMKII-GFP bilaterally into the BLA to selectively target principal neurons. Wireless devices were implanted into the same sites bilaterally two weeks after viral injection (**Fig. 4*A*-*C***). Rats then underwent fear conditioning one week after device implantation (n = 8-9/group; **Fig. 4*A***). Two-way RM ANOVA was conducted for fear conditioning, revealing a significant main effect of trial (*F_5,75_* = 3.06, *P* = 0.01), but no main effect of virus (*F_1,15_* = 0.27, *P* = 0.61) or interaction of virus × trial (*F_5, 75_* = 0.48, *P* = 0.79) (**Fig. 4*D***). Rats then underwent cued fear retrieval with red μLED channel turned on or off in a two-day testing procedure (counterbalanced). Two-way RM ANOVA revealed a significant main effect of μLED (*F_1, 15_* = 4.37, *P* = 0.05) and an interaction of μLED × virus (*F_5, 75_* = 0.48, *P* = 0.79), but no significant main effect of virus (*F_1, 15_* = 3.16, *P* = 0.10). Post hoc analysis showed that μLED illumination reduced freezing in Jaws rats (*t_15_* = 3.78, *P* = 0.004) but not in control GFP rats (*t_15_* = 0.69, *P* = 0.997; **Fig. 4*E***).

**Fig. 4.**
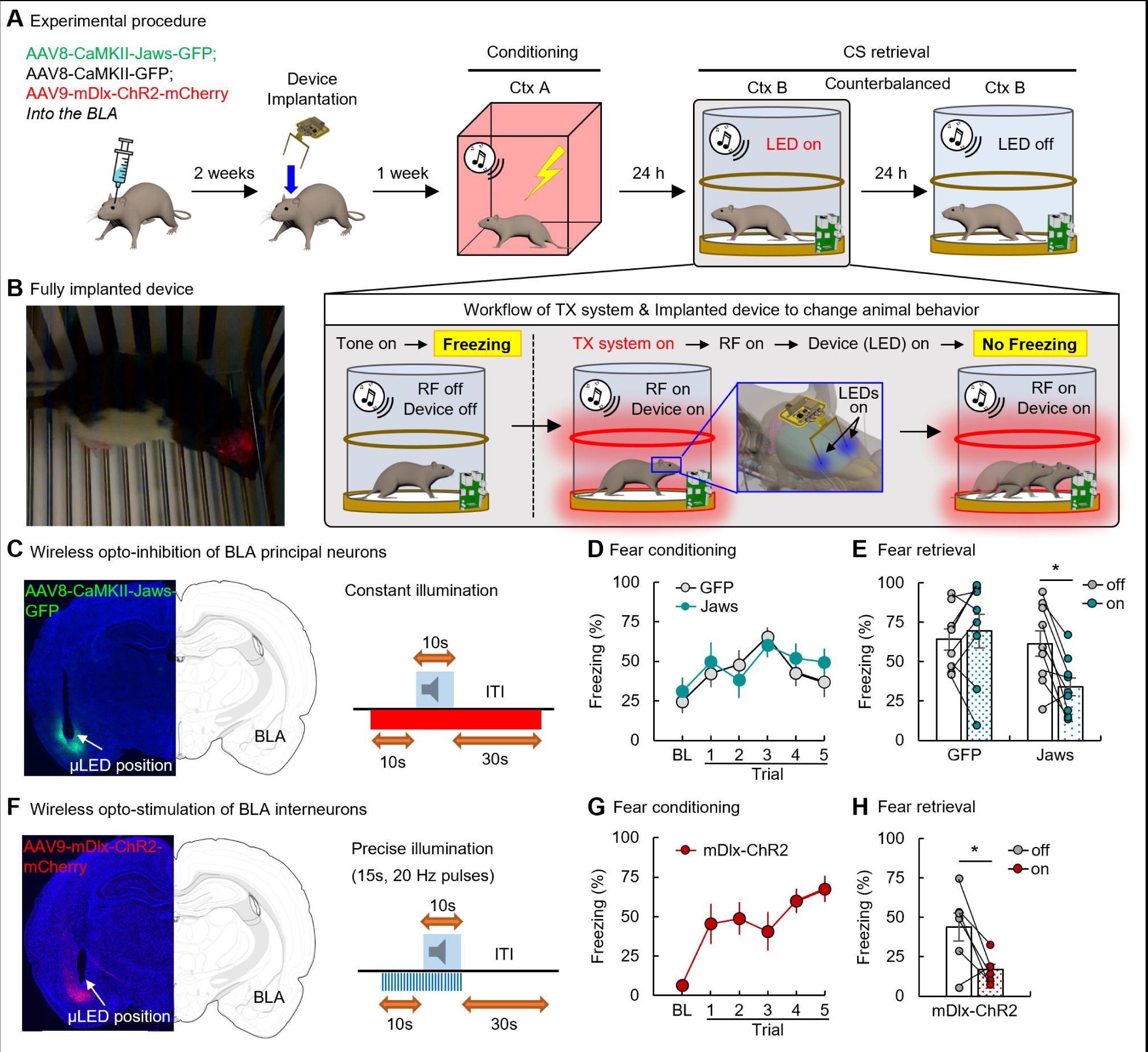
Wireless optogenetic modulation of BLA activity reduces fear. (A) Schematic of experimental procedure and wireless optogenetics. (B) A representative image showing a rat with implanted wireless device (red light indicates that the device is turned on). (C) A representative image of CaMKII-Jaws expression and placement of device implantation in the BLA and the constant opto-stimulation paradigm. (D) Fear conditioning of rats expressing Jaws (n=8) or GFP (n = 9) in BLA principal neurons. (E) Effects of opto-inhibition of BLA principal neurons on fear retrieval. (F) Representative image of mDlx-ChR2 expression and placement of device implantation in the BLA and the precise opto-stimulation paradigm. (G) Fear conditioning of rats expressing ChR2 in BLA interneurons (n = 6). (H) Effects of opto-stimulation of BLA interneurons on fear retrieval. Data are shown as mean ± SEM. *p < 0.05. BLA, basolateral amygdala; CS, conditioned stimulus.

Besides principal neurons, interneurons in the BLA have also been implicated in regulating fear retrieval(26). Interneurons exhibit specific gene expression compared to principal neurons. For example, mDlx has been shown to express in interneurons predominantly(27). In separate experiments, we further validated the TX system by targeting BLA interneurons with the virus AAV9-mDlx-ChR2-mCherry-Fishell 3 (**Fig. 4*F***) and delivering blue light to the BLA to activate these inhibitory cells. Rats underwent the same procedure as described above (n = 6/group; **Fig. 4*A***). One-way RM ANOVA showed a significant main effect (*F_5, 30_* = 4.02, *P* = 0.007), indicating that rats were conditioned successfully (**Fig. 4*F* and *G***). The retrieval data analyzed by a two-tailed paired *t*-test revealed that blue μLED illumination decreased cued fear retrieval (*t_5_* = 2.69, *P* = 0.04; **Fig. 4*H***). Taken together, these data showed that wireless optogenetic manipulation of BLA activity using two different wavelengths of light targeting distinct cellular targets can disrupt cued fear retrieval, validating the TX system in freely moving animals (***SI Appendix*, Movie S3**).

## Discussion

In a recent study, we introduced a software based on custom-trained DeepLabCut for real-time detection of snout-tail pairs of multiple mice in video frames(20, 28). In that study, we succeeded in optimally controlling the TX coil antenna for efficient wireless power delivery by estimating the movement direction of multiple mice. However, this approach was able to determine only the instantaneous directionality of the subject and it was difficult to analyze detailed mouse behaviors with this approach. Here, in this study, we developed AI-driven software capable of detecting specific behaviors (e.g., freezing behavior) of multiple animals by identifying and localizing multiple body parts using DeepLabCut, a CNN variant. With this, we could readily calibrate the sensitivity to detect freezing behavior by changing the thresholds of movement defined by observational studies of freezing behavior. In addition, because this AI-enhanced automated video analysis allows for the analysis of specific behaviors in multiple animals in real-time, scalability to additional behavioral experiments can be expected.

Recently, we developed a low-power actuation mechanism for wireless dual-channel operation(18, 19, 29). The Reed switch-enabled switching mechanism allows optogenetic modulation of neural activity in a freely behaving animal: one channel for stimulation and the other for inhibition. Similar platforms have been proposed that can provide an operational mode of stimulation across both channels. However, these approaches only allow for non-reversible, one-time switching between channels or suffer from poor channel retention (e.g., unexpected switching by environmental electromagnetic noise). In this study, we proposed a new switching mechanism, capacitor-assisted channel switching. Here, a capacitor with a capacitance of 10 μF is connected with a reed switch in parallel. The increased time constant makes the switching circuit less sensitive to unexpected short electromagnetic pulse waves from the surroundings or glitches associated with signal delays. The parallel combination of a capacitor and a reed switch was found to ensure robust channel switching operation.

Compared to traditional tethered optogenetics, wireless optogenetics shows a considerable advantage due to the absence of a tether between the experimental subjects and the optogenetic equipment(30). Our present study validated the application of the TX system-based wireless optogenetics using a classical fear conditioning model, a widely used behavioral task depending on emotional memory. Our data show that wireless optogenetic modulation of BLA neurons disrupted fear retrieval. Notably, selective optical modulation of BLA principal neurons or interneurons via controlling matched μLEDs of the TX system-based dual-channel wireless optogenetic device achieved the same behavioral outcome. These results demonstrated the utility of the TX system-based dual-channel wireless optogenetics in freely moving animals. The greatest advantage of our platform is to offer the opportunity to simultaneously manipulate different types of neurons or brain areas of a group of freely moving animals. The application of this system will advance our understanding of the neurobiological mechanisms underlying complex behaviors and brain disorders that cannot be achieved by traditional optogenetics. For example, it can be used to study functional and causal interactions between different subgroups of neurons either in a local brain area or brain circuits in social fear learning(31). Furthermore, with added wireless recording technology (e.g., wireless electrophysiology or photodiode) the TX system-based wireless device can be designed as a closed-loop system, which will tremendously extend its application in behavioral neuroscience.

## Methods

### Body part detection and evaluation

Complete methods are described in *SI Appendix*, Supplementary text.

### Freezing behavior analysis

Complete methods are described in *SI Appendix*, Supplementary text.

### Quantitative performance assessment of the behavior analysis algorithm

Complete methods are described in *SI Appendix*, Supplementary text.

### Device fabrication

Complete methods are described in *SI Appendix*, Supplementary text.

### TX system setup

Complete methods are described in *SI Appendix*, Supplementary text.

### Thermal characteristics of the implant device

Complete methods are described in *SI Appendix*, Supplementary text.

### Animals

Complete methods are described in *SI Appendix*, Supplementary text.

### Viruses

Complete methods are described in *SI Appendix*, Supplementary text.

### Surgeries and Viruses

Complete methods are described in *SI Appendix*, Supplementary text.

### Behavioral Apparatus

Complete methods are described in *SI Appendix*, Supplementary text.

### Behavioral Procedures and Wireless Optogenetics

Complete methods are described in *SI Appendix*, Supplementary text.

### Statistics

Complete methods are described in *SI Appendix*, Supplementary text.

## Code and materials availability

The main data supporting the results in this study are available within the paper and its Supplementary Information. The code is available from Github (https://github.com/Turmac/freezing_behavior_analysis.git).

## Acknowledgments

This work was supported by grants from the interdisciplinary X-Grants Program, part of the President’s Excellence Fund at Texas A&M University (B.Y., J.M., S.M., S.P.), 2018 NARSARD Young Investigator Awards (S.P.) from Brain & Behavior Research Foundation and National Science Foundation Engineering Research Center for Precise Advanced Technologies and Health Systems for Underserved Populations PATHS-UP (EEC-1648451; S.P.). S.P. would like to express thanks to Dr. Gerald Coté (Texas A&M University) for general advice.

## Author contributions

W.S.K., J.L., Q.L., S.M., and S.P. designed research; W.S.K., J.L., Q.L., S.H., K.Q., R.C, J.M. performed research; W.S.K., J.L., Q.L, S.H., B.Y., S.M., and S.P. analyzed data; and W.S.K., J.L., Q.L., S.M., and S.P wrote the paper.

## References

1. P. Rajasethupathy, E. Ferenczi, K. Deisseroth, Targeting Neural Circuits. Cell 165, 524–;534 (2016).

2. K. Deisseroth, Optogenetics. Nat. Methods 8, 26–;29 (2011).

3. K. Deisseroth, Optogenetics: 10 years of microbial opsins in neuroscience. Nat. Neurosci. 18, 1213–;1225 (2015).

4. L. Grosenick, J. H. Marshel, K. Deisseroth, Closed-loop and activity-guided optogenetic control. Neuron 86, 106–;139 (2015).

5. A. L. Halberstadt, Automated detection of the head-twitch response using wavelet scalograms and a deep convolutional neural network. Sci. Rep. 10, 8344 (2020).

6. A. Krizhevsky, I. Sutskever, G. E. Hinton, ImageNet classification with deep convolutional neural networks. Commun. ACM 60, 84–;90 (2017).

7. K. He, G. Gkioxari, P. Dollar, R. Girshick, Mask R-CNN in 2017 IEEE International Conference on Computer Vision (ICCV), (IEEE, 2017), pp. 2980–;2988.

8. L. C. Chen, G. Papandreou, I. Kokkinos, K. Murphy, A. L. Yuille, DeepLab: Semantic Image Segmentation with Deep Convolutional Nets, Atrous Convolution, and Fully Connected CRFs. IEEE Trans. Pattern Anal. Mach. Intell. 40, 834–;848 (2018).

9. A. Mathis, et al., DeepLabCut: markerless pose estimation of user-defined body parts with deep learning. Nat. Neurosci. 21, 1281–;1289 (2018).

10. M. Danielczuk, et al., Segmenting Unknown 3D Objects from Real Depth Images using Mask R-CNN Trained on Synthetic Data in 2019 International Conference on Robotics and Automation (ICRA), (IEEE, 2019), pp. 7283–;7290.

11. L. John, M. Andrew, C. N. P. Fernando, Conditional Random Fields: Probabilistic Models for Segmenting and Labeling Sequence Data. ICML ‘01 Proc. Eighteenth Int. Conf. Mach. Learn. 2001, 282–;289 (2001).

12. L. Fenno, O. Yizhar, K. Deisseroth, The Development and Application of Optogenetics. Annu. Rev. Neurosci. 34, 389–;412 (2011).

13. O. Yizhar, L. E. Fenno, T. J. Davidson, M. Mogri, K. Deisseroth, Optogenetics in Neural Systems. Neuron 71, 9–;34 (2011).

14. P. Bonnavion, A. C. Jackson, M. E. Carter, L. de Lecea, Antagonistic interplay between hypocretin and leptin in the lateral hypothalamus regulates stress responses. Nat. Commun. 6, 6266 (2015).

15. F. Pisanello, et al., Multipoint-emitting optical fibers for spatially addressable in vivo optogenetics. Neuron 82, 1245–;54 (2014).

16. M. A. Parent, L. M. Amarante, B. Liu, D. Weikum, M. Laubach, The medial prefrontal cortex is crucial for the maintenance of persistent licking and the expression of incentive contrast. Front. Integr. Neurosci. 9, 1–;14 (2015).

17. A. R. Adamantidis, F. Zhang, A. M. Aravanis, K. Deisseroth, L. De Lecea, Neural substrates of awakening probed with optogenetic control of hypocretin neurons. Nature 450, 420–;424 (2007).

18. W. S. Kim, M. Jeong, S. Hong, B. Lim, S. Il Park, Fully implantable low-power high frequency range optoelectronic devices for dual-channel modulation in the brain. Sensors (Switzerland) 20, 1–;14 (2020).

19. W. S. Kim, et al., Organ-specific, multimodal, wireless optoelectronics for high-throughput phenotyping of peripheral neural pathways. Nat. Commun. 12, 157 (2021).

20. H.-M. Woo, et al., Machine Learning Enabled Adaptive Wireless Power Transmission System for Neuroscience Study in 2020 54th Asilomar Conference on Signals, Systems, and Computers, (IEEE, 2020), pp. 808–;812.

21. Jia Deng, et al., ImageNet: A large-scale hierarchical image database. 248–;255 (2009).

22. Z. T. Pennington, et al., ezTrack: An open-source video analysis pipeline for the investigation of animal behavior. Sci. Rep. 9, 1–;11 (2019).

23. M. J. Caruso, T. Bratland, C. H. Smith, R. Schneider, A new perspective on magnetic field sensing. Sensors (Peterborough, NH) 15, 34–;46 (1998).

24. J. Mankowski, M. Kristiansen, A review of short pulse generator technology. IEEE Trans. Plasma Sci. 28, 102–;108 (2000).

25. S. Maren, Neurobiology of Pavlovian Fear Conditioning. Annu. Rev. Neurosci. 24, 897–;931 (2001).

26. M. B. Perumal, P. Sah, Inhibitory Circuits in the Basolateral Amygdala in Aversive Learning and Memory. Front. Neural Circuits 15, 1–;7 (2021).

27. J. Dimidschstein, et al., A viral strategy for targeting and manipulating interneurons across vertebrate species. Nat. Neurosci. 19, 1743–;1749 (2016).

28. W. S. Kim, et al., AI-enabled, implantable, multichannel wireless telemetry for photodynamic theraphy. 1–;21 (2021).

29. W. S. Kim, et al., A soft, biocompatible magnetic field enabled wireless surgical lighting patty for neurosurgery. Appl. Sci. 10 (2020).

30. S. Il Park, et al., Soft, stretchable, fully implantable miniaturized optoelectronic systems for wireless optogenetics. Nat. Biotechnol. 33, 1280–;1286 (2015).

31. J. Debiec, A. Olsson, Social Fear Learning: from Animal Models to Human Function. Trends Cogn. Sci. 21, 546–;555 (2017).

