## Supplementary Information for "AI-driven high-throughput automation of behavioral analysis and dual-channel wireless optogenetics for multiplexing brain dynamics"

**This PDF file includes:**

Supplementary text
Figures S1 to S7
Table S1
Legends for Movies S1 to S3
SI References

**Other supplementary materials for this manuscript include the following:**

Movies S1 to S3

**Supplementary Information Text**

**Methods**

**Body part detection and evaluation**

Mathis et al. investigated the body part detection task where the camera was placed on the above the mice and the goal was to detect the snout, left ear, right ear, and tail(1). Compared with Mathis et al.'s task, our task is more challenging in the following aspects: (1) in our task the camera was placed closely in front of the mice (side view) so that there was a higher variety of body poses; (2) instead of only 4 body parts, our goal was to detect a total of 9 body parts. Therefore, in order to effectively train the network, we manually labeled 475 frames out of 5 videos (>54,000 frames) that DeepLabCut used. We randomly split the labeled dataset into a training set with 451 frames (95%) and a test set with 24 frames (5%). We chose the ResNet-50(2) pre-trained on the ImageNet(3) dataset. We trained the network with the SGD (Stochastic Gradient Descent) optimizer and a batch size of 1 for 300k iterations. We performed both the training and testing with an NVIDIA GTX 1080 Ti GPU with 11G memory.

**Freezing behavior analysis**

We ran the trained network on all the video frames to detect the body parts. In addition to the position of each body part, the detection network also outputs the likelihood of the detection. We set a cutoff threshold $T_1$ for the likelihood, such that the detection is invalid if the likelihood is less than $T_1$. With all the detected body part positions, we defined two metrics to measure the motion of the subject over a short period of time T: $d_s$ and $d_c$. The $d_s$ measured the total moving distance of a specific body part, while $d_c$ measured the total distance from the center of the body part during T. We took the mean value of all the detected body parts for both $d_s$ and $d_c$, since not all the body parts were always visible. Finally, we defined two thresholds $T_s$ and $T_c$ according to the experimental condition, to decide whether the subject was freezing or not. We analyzed the freezing behavior for every frame in a sliding window manner.

**Quantitative performance assessment of the behavior analysis algorithm**

Most currently available freezing analysis methods are based on pixel differences of the video frames, e.g., ezTrack(4). To demonstrate the effectiveness of our proposed freezing behavior analysis method, we compared our system with ezTrack and hand-scoring. We captured 4 test videos and asked three human experts to label the freezing behavior for all four test videos. We then compared our result and ezTrack's result with hand scoring and reported the difference in number of seconds per minute. We also announced the average variability among three human experts as the baseline.

**Device fabrication**

The device's pattern fabrication process started with attaching a thin, flexible copper/polyimide (Cu/PI) bilayer film (thickness; 12 μm/18 μm, AC181200RY, DupontTM Pyralux®) onto a glass slide (dimensions, 5.08 cm by 7.62 cm). In a cleanroom, we deposited the photoresistor (AZ 1518, AZ®) onto the prepared substrate for 2 μm thick (recipe; spin-coated at 3,000 r.p.m. for 30 sec), and illuminated UV lights to lithograph (EVG610, EV Group) patterns (recipe; UV intensity for 120 $mJ/cm^2$). In order to carve the photoresistor layer, the sample was submerged in developer solution (AZ Developer 1:1, AZ®) for 1 min. Next, the sample was immersed in copper etchant (LOT: Z03E099, Alfa AesarTM) until patterns were well-etched and the sample was rinsed with solvents (acetone, methanol, and isopropanol successively) and distilled water for 1 min. After fully drying the samples, in the standard laboratory, chip components including SMD (surface-mount-device) LEDs, passive components, and IC components were mounted onto the pattern using a soldering machine by hand. Finally, we applied Polydimethylsiloxane (PDMS) (SylgardTM 184 silicone elastomer kit, Dow®; 10:1 mix ratio) with a dip-coating process (500 μm thick) to the sample, and then baked it in an oven at 100 ºC for 90 min for encapsulation.

**TX system setup**

We fabricated a customized antenna with 8-ga bare Cu wire for the secondary-antenna coil and Cu stripes (0.635 mm thick by 2.54 cm wide) for the primary-antenna coil. The secondary-coils were wrapped around the center-height of the cage. Impedance matching using Network Analyzer (ENA Series E5063A, Keysight) with a discrete capacitor component yielded primary- and secondary-coils, each of which resonated at 13.56 MHz (the primary-stripes) and 15 MHz (the secondary-coils), respectively. Wireless power control systems consisted of an RF power generator (ID ISC.LRM2500-A, FEIG Electronics), and an auto-tunable matching board (ID ISC.DAT-A, FEIG Electronics). The multi-cage system required a TX controller including an RF multiplexer (ID ISC.ANT.MUX.M8, FEIG Electronics), and a control board (nRF52832 Development Kit, Nordic semiconductor) that has been reprogrammed from ver.13.0.0 (nRF52 SDK 13.0.0.0, Nordic semiconductor).

**Thermal characteristics of the implant device**

We measured thermal variations of the light sources in the implant device using an infrared camera (VarioCAM HDx head 600, InfraTech) in two configurations: a device installed under a sealed bag of saline solution (10 % PBS) instead of a rat, and a device itself in the cage. The power supply was provided constantly for 20 mins, the trickiest case for the device's operation in temperature changes.

**Animals**

Adult naïve male and female Long-Evans Blue Spruce rats (200 - 240 g upon arrival) were obtained from Envigo (Indianapolis, IN). Rats were individually housed in clear plastic cages on rotating racks in a climate-controlled vivarium with a fixed 14:10 hour light: dark cycle (lights on at 7:00 AM). All experiments were conducted during the light phase. Rats were given access to standard rodent chow and water ad libitum. Upon arrival, all rats were handled by the experimenter (~30 sec/rat/day) for a minimum of 5 days prior to the start of any surgical or behavioral procedures. All experimental procedures were conducted in accordance with the US National Institutes of Health (NIH) Guide for the Care and Use of Laboratory Animals and were approved by the Texas A&M University Institutional Animals Care and Use Committee (IACUC).

**Viruses**

AAV8-CaMKII-Jaws-KGC-GFP-ER2-WPRE-SV40 (AAV8-CaMKII-Jaws-GFP) was purchased from University of Pennsylvania Vector Core. AAV8-CaMKII-GFP was purchased from UNC vector core. AAV9-mDlx-ChR2-mCherry was purchased from Addgene (Watertown, MA). Viruses were diluted to particular titers with sterilized 1x DPBS. The final titers of viruses injected into the BLA were $5.8 \times 10^{12}$ GC/mL for AAV8-CaMKII-Jaws-GFP, $4.2 \times 10^{12}$ GC/mL for AAV8-CaMKII-GFP, and $8.0 \times 10^{12}$ GC/mL for AAV9-mDlx-ChR2-mCherry.

**Surgeries and Viruses**

Rats were anesthetized with isoflurane (5% for induction, 1-2% for maintenance) and placed into a stereotaxic frame (Kopf Instruments). The hair on the scalp was shaved, povidine-iodine was applied to the skin, and a small incision was made in the scalp to expose the top of the skull. The skull was leveled by placing bregma and lambda in the same horizontal plane. Small holes were drilled in the skull to affix four jeweler's screws.

For viral injection, rats received bilateral infusions of viruses into the BLA (0.5 µl/side; AAV8-CaMKII-Jaws-GFP, AAV8-CaMKII-GFP, or AAV9-mDlx-ChR2-mCherry). Virus was infused at a rate of 0.1 µL/min and injector tips were left in the brain for five additional minutes to allow for diffusion. The coordinates for BLA viral injection were: AP: -2.85 mm, ML: -4.85 mm, DV: -8.7 mm (relative to bregma surface). Two weeks after viral injection, probes of wireless devices were implanted into the BLA bilaterally (same coordinate as viral injection). Dental cement was used to secure the wireless device to the skull. Topical antibiotic (Triple Antibiotic Plus; G&W Laboratories) was applied to the surgical site and one chewable carprofen tablet (2 mg; Bio-Serv) was provided for post-operative pain management. Rats were given one week after device implantation for appropriate viral expression (3 weeks) and to recover prior to the beginning of behavioral testing.

**Behavioral Apparatus**

Fear conditioning was conducted in 8 identical rodent fear conditioning chambers (context A; 30 × 24 × 21 cm; Med Associates). Each chamber consisted of two aluminum sidewalls and a Plexiglas ceiling and rear wall, a hinged Plexiglas door, and a grid floor. The grid floor consisted of 19 stainless steel rods that were wired to a shock source and a solid-state grid scrambler for delivery of the footshocks (Med Associates). A speaker for delivering auditory stimuli, ventilation fans, and house lights were installed in each chamber. Each conditioning chamber rested on a load-cell platform that was used to record chamber displacement in response to each rat's motor activity and was acquired online via Threshold Activity software (Med-Associates). Freezing was quantified by computing the number of observations for each rat that had a value less than the freezing threshold (load-cell activity = 10).

Fear retrieval was conducted in a custom-made clear plastic cylinder (context B; 30 × 45 cm; D × H) with a clear plastic movable cover. The cylinder was covered by black and white wallpapers to provide a visual cue distinct with the conditioning chamber. The antenna of the TX system was equipped around the outer surface of the cylinder. The cylinder was placed in a room different from where the conditioning chambers are located. The house light of the room was kept on during experimentation to provide enough light for videotaping. An iPad was placed near the top of the cylinder to provide auditory stimuli (80 dB, 2 kHz). The behavior was videotaped by a cell phone placed on the cylinder cover. The cylinder was cleaned with 70 % ethanol prior to each behavioral session. Videos and freezing behaviors were analyzed using the open-source tool ezTrack(4).

**Behavioral Procedures and Wireless Optogenetics**

An overview of the behavioral experiments is provided in Figure 4a. Rats were implanted with wireless devices two weeks after viral injection. One week after device implantation, rats underwent auditory fear conditioning. Fear conditioning was conducted in context A [3-min BL, five tones (10 s, 80 dB, 8 kHz)-footshock (US; 1.0 mA, 2 s) pairings with 60 s intertrial intervals (ITIs), and an additional 60 s post-shock period]. Fear retrieval [3-min BL; 5 × 10 s tones; 30 s ITIs] was conducted in context B (a plastic cylinder equipped with TX system) 1-2 days after conditioning with a counterbalanced design of μLED on and off.

The rat expressing Jaws or control GFP in BLA principal neurons was bilaterally illuminated using the red μLED channel of the wireless device (μLED was turned on 10 s before the first tone onset and persisted to the end of testing). The rat expressing mDlx-ChR2 in BLA interneurons was bilaterally illuminated using the blue μLED channel of the wireless device (μLED was turned on 10 s before each tone onset and turned off at each tone offset).

Upon completion of the experiment, rats were overdosed with sodium pentobarbital (Fatal Plus; 100 mg/mL, 0.5 mL, i.p.) and perfused transcardially with physiological saline followed by 10% formalin. Brains were extracted and stored about 16-18 h (at 4° C) in 10 % formalin after which they were transferred to a 30 % sucrose solution for a minimum of 5 days. Brains were then sliced using a cryostat (Leica Microsystems) at -20° C. Viral expression was verified with a Zeiss microscope (Axio Imager).

**Statistics**

All data were represented as means ± SEM. Data were analyzed using Prism GraphPad 9.0. For conditioning, two-way or one-way repeated-measures (RM) analysis of variance (ANOVA) was conducted. For the retrieval in CaMKII-Jaws experiment, two-way RM ANOVA with laser and memory age as within-subjects factors and virus as between-subjects factor was conducted. For the retrieval in mDlx-ChR2 experiment, two-tailed paired t-test was conducted. Significant ANOVA was followed by post hoc Bonferroni's multiple comparisons test. Group sizes were determined based on prior work and what is common in the field. $P < 0.05$ was considered statistically significant.

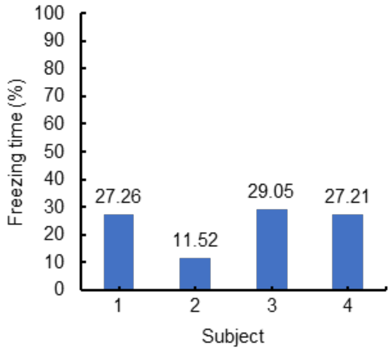


**Fig. S1.** Statistics of freezing analysis with unconditioned animals. The average percentage of no movements in natural behaving rats is 23.88 %.

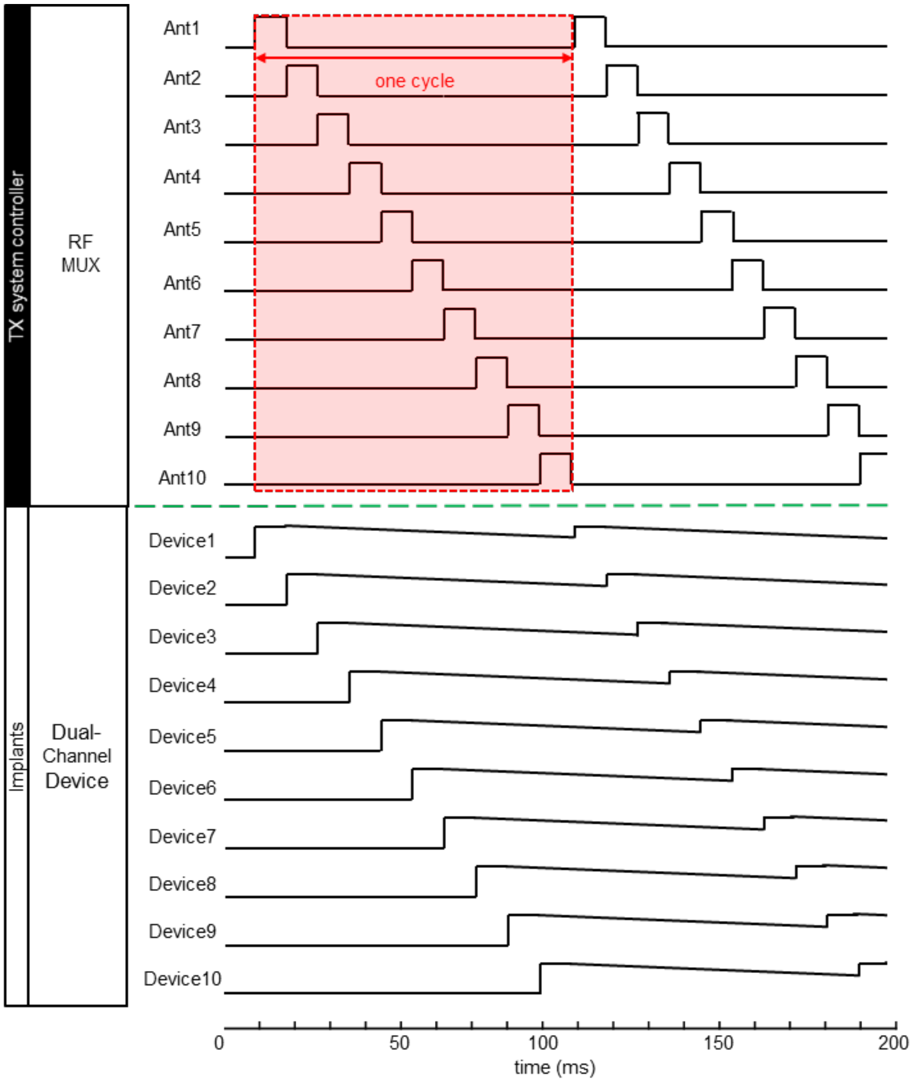


**Fig. S2.** Time division multiplexing scheme and representative waveforms at each antenna and implant. Assume the duty cycle 10% with 10 ms pulse train.

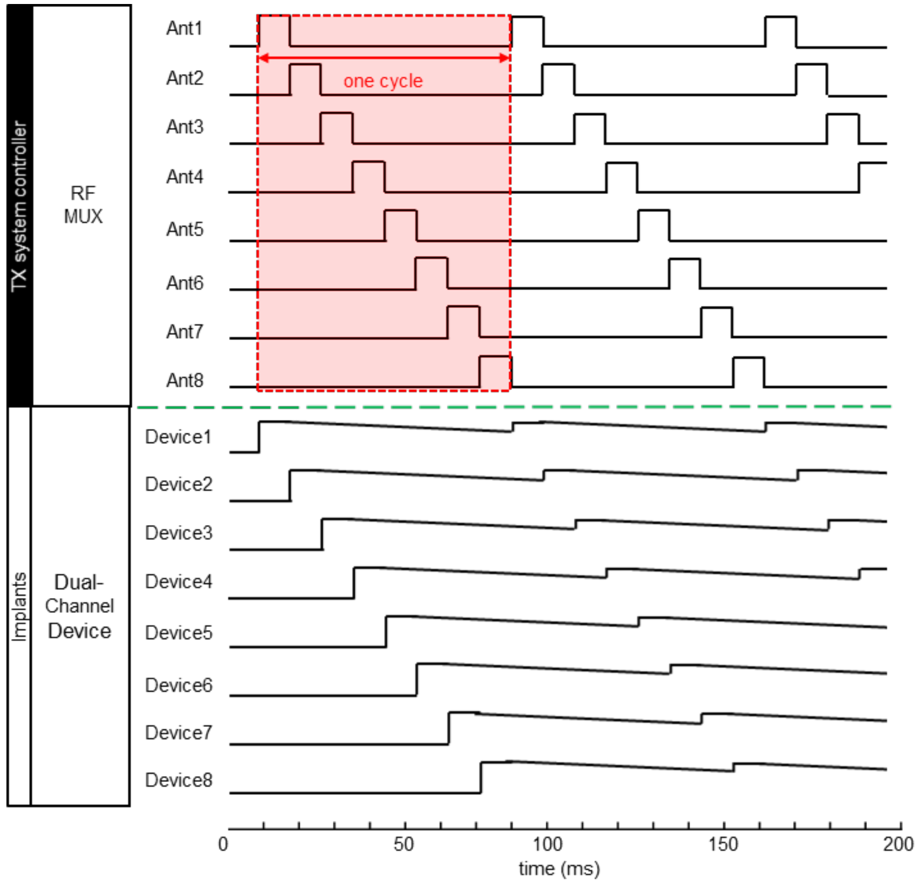


**Fig. S3.** Time division multiplexing scheme and representative waveforms at each antenna and implant. Assume the duty cycle 12.5% with 10 ms pulse train.

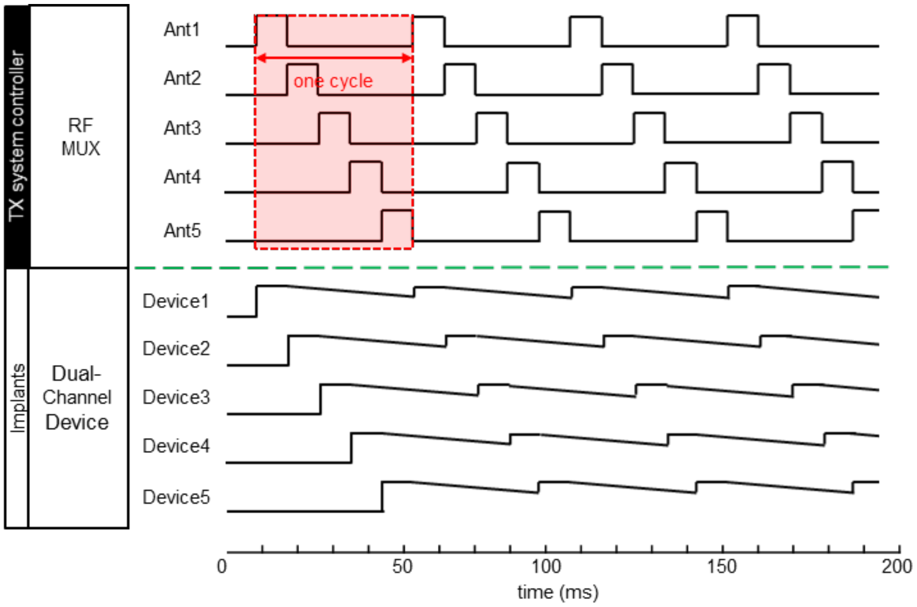


**Fig. S4.** Time division multiplexing scheme and representative waveforms at each antenna and implant. Assume the duty cycle 20% with 10 ms pulse train.

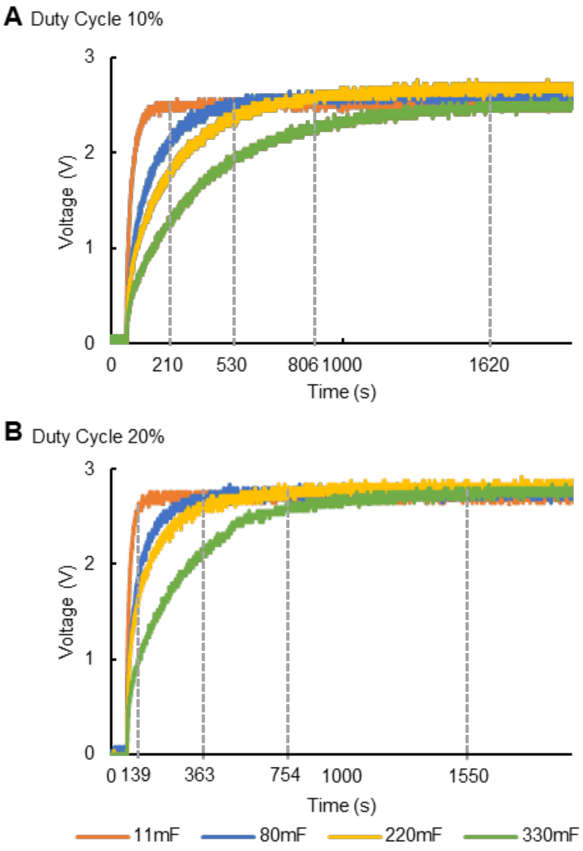


**Fig. S5.** Plot of threshold for switching as a function of capacitance according to duty cycles with 10ms pulse train: (*A*) 10%, (*B*) 20%.

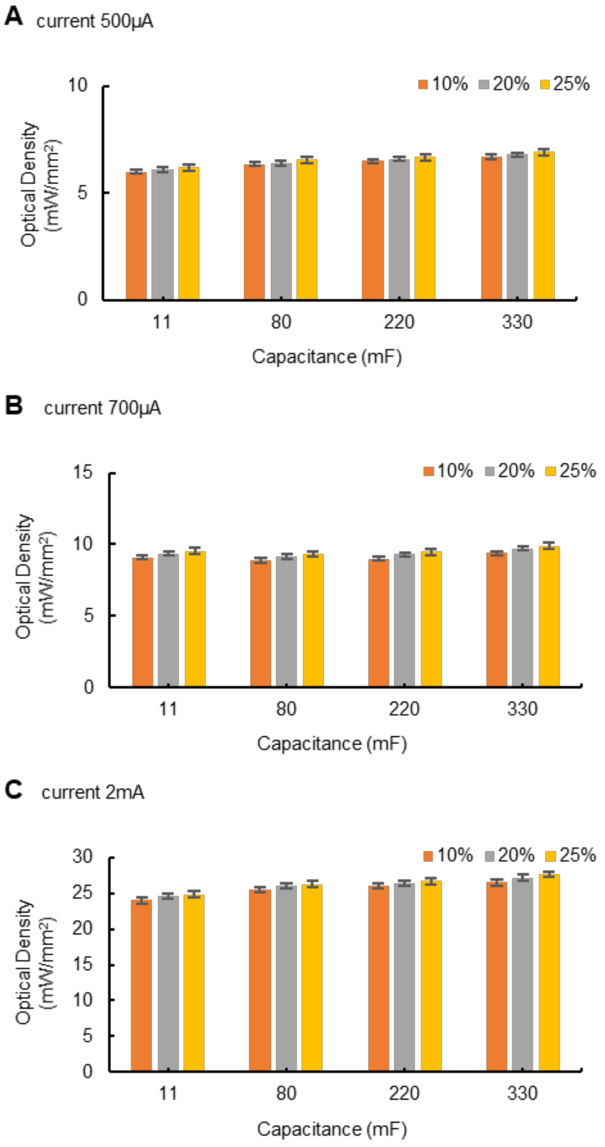


**Fig. S6.** Optical intensity vs. capacitance as a function of duty cycle with each different current: (*A*) 500 µA, (*B*) 700 µA, and (*C*) 2 mA.

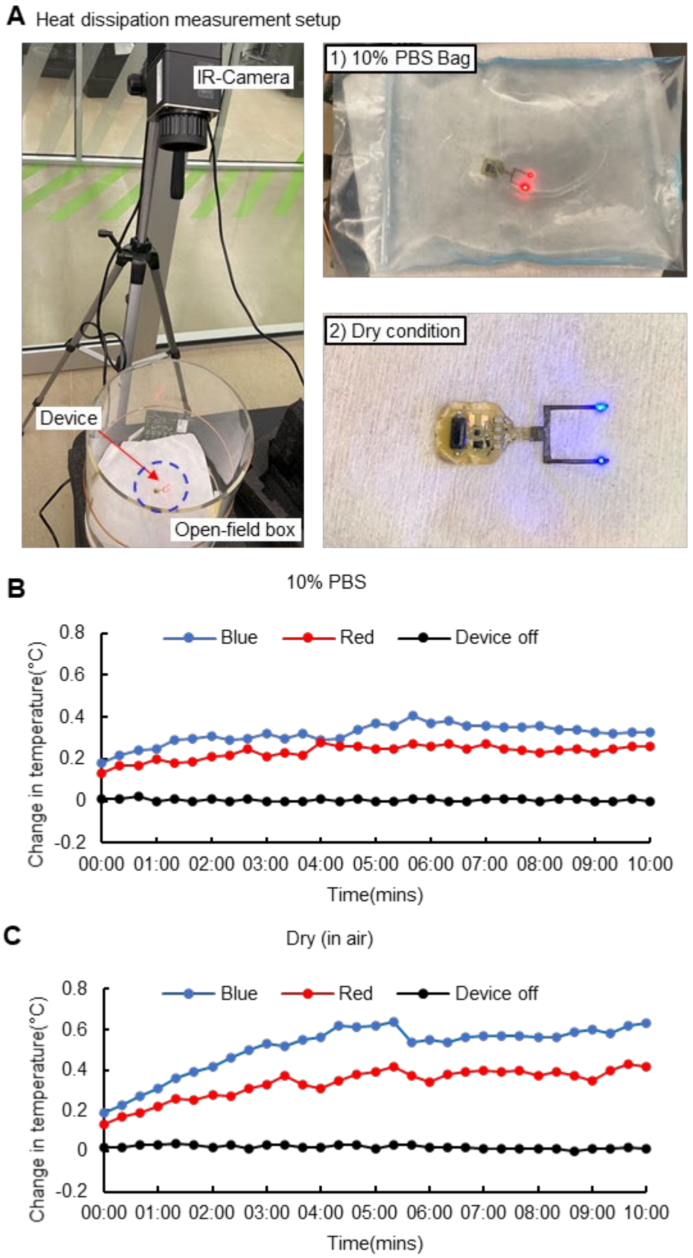


**Fig. S7.** (*A*) Pictures of an experimental setup for wireless measurements of heat dissipation using IR camera (left). Here, TX power is set to 4 W. The two right images show a device mounted on sealed bag of 10 % PBS saline solution (right to the top), and itself in a cage (right to the bottom), respectively. Plots of optical intensity as a function of constant time in each condition; 10 % PBS bag (*B*), and dry (*C*).

| **Package name** | **Version** |
|---|---|
| altair | 4.1.0 |
| deeplabcut | 2.1.10.4 |
| matplotlib | 3.1.3 |
| numpy | 1.17.5 |
| opencv-python | 4.5.2.54 |
| pandas | 1.3.0 |
| Pillow | 8.3.1 |
| tensorflow-gpu | 1.15.0 |

**Table S1.** Summary of resulting python package information.

**Movie S1 (separate file).** *In vivo* demonstration of multiple-animals analysis in multiple cages.

**Movie S2 (separate file).** Demonstration of switching mechanisms.

**Movie S3 (separate file).** *In vivo* demonstration of freezing behavior disruption by optogenetics.

**SI References**


1. A. Mathis, *et al.*, DeepLabCut: markerless pose estimation of user-defined body parts with deep learning. *Nat. Neurosci.* **21**, 1281–1289 (2018).
2. Jia Deng, *et al.*, ImageNet: A large-scale hierarchical image database. 248–255 (2009).
3. A. Krizhevsky, I. Sutskever, G. E. Hinton, ImageNet classification with deep convolutional neural networks. *Commun. ACM* **60**, 84–90 (2017).
4. Z. T. Pennington, *et al.*, ezTrack: An open-source video analysis pipeline for the investigation of animal behavior. *Sci. Rep.* **9**, 1–11 (2019).
